# The impact of cell geometry and the cytoskeleton on the nucleo-cytoplasmic localisation of the SMYD3 methyltransferase suggests that the epigenetic machinery is mechanosensitive

**DOI:** 10.1101/2020.03.17.994046

**Authors:** David Pereira, Alain Richert, Souhila Medjkane, Sylvie Hénon, Jonathan B Weitzman

**Affiliations:** Université de Paris, Matière et Systèmes Complexes, CNRS UMR 7057, Paris, France; Université de Paris, Epigenetics and Cell Fate, CNRS UMR 7216, Paris, France

## Abstract

Mechanical cues from the cellular microenvironment are converted into biochemical signals controlling diverse cell behaviours, including growth and differentiation. But it is still unclear how mechanotransduction ultimately affects nuclear readouts, genome function and transcriptional programs. Key signaling pathways and transcription factors can be activated, and can relocalize to the nucleus, upon mechanosensing. Here, we tested the hypothesis that epigenetic regulators, such as methyltransferase enzymes, might also contribute to mechanotransduction. We found that the SMYD3 lysine methyltransferase is spatially redistributed dependent on cell geometry (cell shape and aspect ratio) in murine myoblasts. Specifically, elongated rectangles were less permissive than square shapes to SMYD3 nuclear accumulation, via reduced nuclear import. Notably, SMYD3 has both nuclear and cytoplasmic substrates. The distribution of SMYD3 in response to cell geometry correlated with cytoplasmic and nuclear lysine tri-methylation (Kme3) levels, but not Kme2. Moreover, drugs targeting cytoskeletal acto-myosin induced nuclear accumulation of Smyd3. We also observed that square vs rectangular geometry impacted the nuclear-cytoplasmic relocalisation of several mechano-sensitive proteins, notably YAP/TAZ proteins and the SETDB1 methyltransferase. Thus, mechanical cues from cellular geometric shapes are transduced by a combination of transcription factors and epigenetic regulators shuttling between the cell nucleus and cytoplasm.

## Introduction

Mechanosensing is a fundamental property of many living cells. As part of their normal physiological functions, cells in multicellular organisms must respond and adapt to mechanical stimuli such as forces, deformations, geometry and stiffness of the extracellular matrix [^1^,^2^, ^3^, ^4^]. Conversely, aberrant mechanical responsiveness is often associated with severe diseases, including cardiovascular disorders, myopathies, fibrotic diseases, or cancer metastasis. Cells can sense and respond to changes in rigidity by aligning their shape, cytoskeletal structures, and traction forces [^5^, ^4^]. For example, the sensing of matrix substrate rigidity and geometrical constraints can direct stem cell lineage specification and differentiation programs [^6^, ^7^, ^2^]. In recent years, significant progress has been made in understanding the signaling pathways of mechanotransduction, and the resulting gene expression regulation by the modulation of transcription factors activity. Notable examples include the transcriptional coactivators, YAP and TAZ [^8^], and MRTF-A [^9^], which are translocated to the nucleus upon mechano-sensitive activation of the Hippo and Rho/MRTF pathways, respectively. Despite these advances, less attention has been paid to modifications of the chromatin and the epigenetic machinery by mechanical cues. Epigenetic regulators could modulate the transcriptional activity and gene expression programs in differentiation, offering a long-term temporal memory of transient spatial mechano-signals.

Epigenetic mechanisms regulate gene expression through a series of post-translational modifications [^10^, ^11^, ^12^, ^13^]. Epigenetic enzymes can modify residues in the histone N-terminal tails to impact chromatin structure and regulate transcriptional activity [^13^, ^14^]. Lysine acetylation and lysine methylation have emerged as key regulatory events in epigenetic control of genome function [^12^, ^15^, ^16^, ^17^]. Many enzymes containing a characteristic SET domain are linked to methylation of lysine residues in histone tails [^18^]. The resulting mono-, di- or tri-methylated lysine residues (Kme1, Kme2 or Kme3) can exert either activating or repressive effects on gene expression [^12^, ^19^]. Despite their initial identification as ‘histone methyltransferases’, many of these enzymes can also modify non-histone proteins, including transcription factors such as p53, YAP or β-catenin [^20^, ^21^, ^22^, ^23^]. Intriguingly, some methyltransferases have been recently linked to cytoplasmic, non-histone substrates, suggesting broader roles in regulating cellular states [^24^, ^23^, ^25^].

SMYD3 is a member of the SMYD (SET and MYND domain-containing proteins) lysine methyltransferases that has been implicated in differentiation and cancer progression [^26^, ^27^, ^28^, ^29^, ^24^, ^30^]. SMYD3 was initially identified as a histone H3K4 methyltransferase enzyme upregulated in metastatic cancers [^31^]. Subsequent studies showed that SMYD3 can also methylate H4K5 and H2A.Z.1, depending on the cellular context [^32^, ^33^]. Furthermore, SMYD3 has cytoplasmic, non-histone targets that contribute to tumorigenesis; for example, SMYD3 can methylate lysine residues in the cytoplasmic domain of the VEGF receptor in angiogenesis or the MAP3K2 signaling kinase in Ras-dependent oncogenesis [^34^, ^24^]. Despite these striking examples of differential SMYD3 substrates in the nucleus or the cytoplasm, little is known about what regulates the nuclear-cytoplasmic shuttling of SMYD3. SMYD family proteins have been linked to muscle differentiation in a wide range of studies: for example, we recently showed that SMYD3 is critical for the differentiation program of mammalian muscle differentiation [^35^]. Furthermore, SMYD3 was linked to cardiac and skeletal muscle development in zebrafish [^36^], and cytoplasmic activity of SMYD2 maintains skeletal muscle functions [^37^]. Thus, muscle differentiation could offer an interesting experimental model to explore a link between cell geometry, mechanical cues and epigenetic SMYD proteins. The differentiation process from myoblasts to differentiated myotubes involves extensive changes in cell morphology (notably cell elongation), as well as gene expression and epigenetic modifications [^38^, ^39^]. Muscle cells appear to have both cytoplasmic and nuclear targets for members of the SMYD methyltransferase family: for example, SMYD2 methylates cytoplasmic Hsp90 chaperone in myoblasts to promote the interaction with titin [^37^], whereas the deletion of Smyd1 impaired myoblast differentiation by decreased expression of muscle-specific genes [^40^].

In this study, we used fibronectin-coated adhesive micropatterns to alter cellular geometry (size, shape, and aspect ratio) of murine myoblasts and to investigate the link between cell shape and contractility and the subcellular localisation of the SMYD3 methyltransferase enzyme. Our findings link geometric and mechanical cues to methyltransferase localization and lysine methylation. We uncovered an important link between the geometric constraints and the actomyosin cytoskeleton of myoblast cell and the regulation of SMYD3 subcellular distribution.

## RESULTS

### Cell geometry regulates SMYD3 distribution

The recent discovery that the SMYD3 methyltransferase has both nuclear and cytoplasmic targets that are critical for cell proliferation, migration and differentiation [^24,30,32–35^], raised the intriguing question how cellular states might regulate nuclear vs. cytoplasmic partitioning. We investigated whether cell spreading affected SMYD3 subcellular localization. We plated C2C12 myoblasts on micropatterns of defined sizes and shapes coated with fibronectin and followed SMYD3 localisation using immunofluorescence of endogenous SMYD3 (WT C2C12 cells) or Flag-tagged constructs (C2C12 cells stably expressing SMYD3-HA-Flag fusion protein). Previous reports have linked cell spreading area to nuclear localisation of transcriptional regulators (e.g. YAP/TAZ) [^8^]. Interestingly, we found that SMYD3 nuclear vs cytoplasmic localisation was affected by the shape of the micropattern, rather than the pattern area *per se*.

We noted that SMYD3 was primarily cytoplasmic in C2C12 cells [Figure 1a], as previously reported for other methyltransferases, such as the related SMYD2 enzyme [^37^]. However, cells plated on rectangular micropatterns with an aspect ratio of 1:5 and 1:8 exhibited more cytoplasmic SMYD3 compared to cells plated on square micropatterns. We observed similar results whether we measured the exogenous SMYD3-HA-Flag [Figure 1a-b] or the endogenous SMYD3 protein [Figure S1c-d].

**Figure 1:**
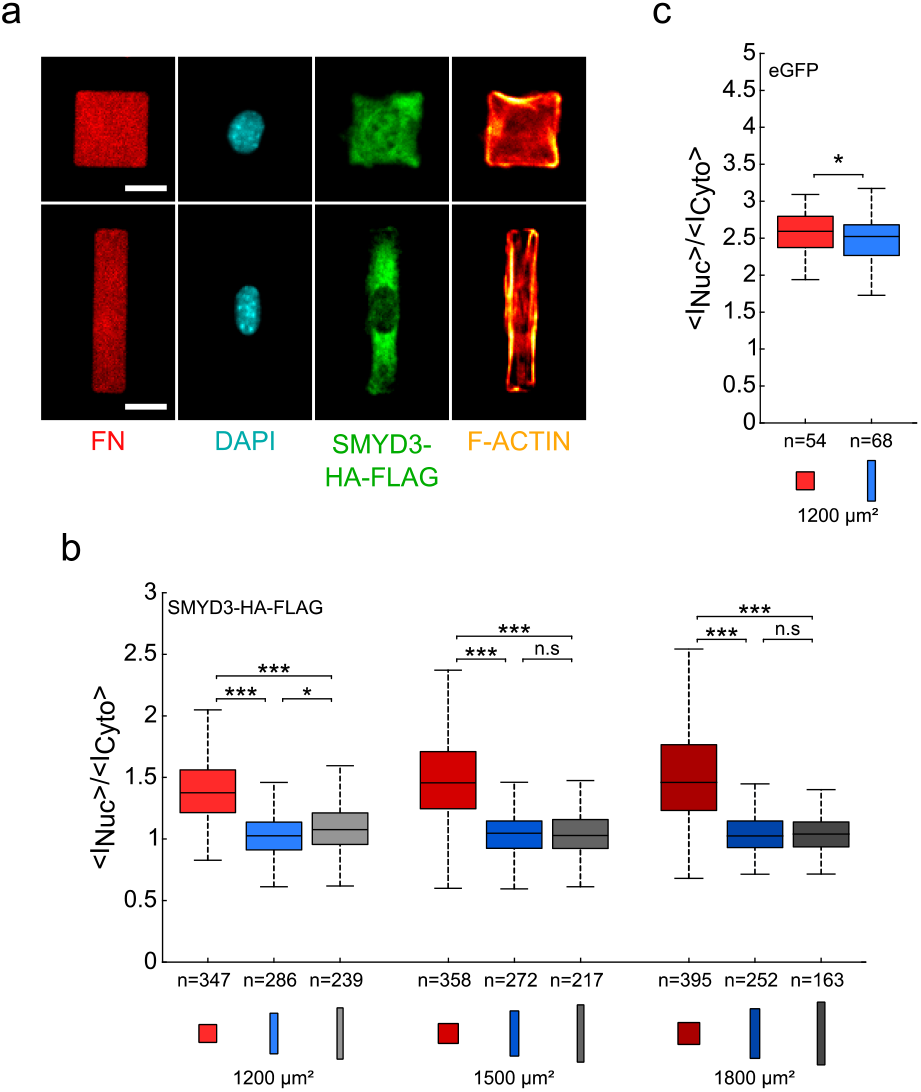
Cell geometry on fibronectin micropatterns regulates SMYD3 distribution. **(a)** Myoblast C2C12 cells expressing SMYD3-HA-FLAG were plated on fibronectin micropatterns with the same area (1200 μm^2^), but different aspect ratios; square (1:1, upper panel) or rectangle (1:5, lower panel). The micrographs show the fibronectin (FN) patterning, the nuclear DNA staining (DAPI), SMYD3-HA-Flag protein location and F-actin (SiR-actin). The SMYD3 nuclear:cytoplasmic distribution appears different between square and rectangle patterns. Scale bars: 20 μm. **(b)** Quantification of the nuclear:cytoplasmic (<I_Nuc_>/<I_Cyto_>) distribution ratio for SMYD3-HA-Flag over a range of pattern areas (1200-1800 μm^2^) and geometries: squares (1:1 aspect ratio, red), rectangles (1:5 aspect ratio, blue) and elongated rectangles (1:8 aspect ratio, grey).The median value of the ratio is 27 to 42% higher for cells plated on squares than on rectangle patterns. **(c)** Quantification of the nuclear:cytoplasmic (<I_Nuc_>/<I_Cyto_>) distribution ratio for control eGFP transfected into C2C12 cells on square or rectangle patterns (1200 μm^2^). The eGFP nuclear:cytoplasmic distribution is slightly different (less than 3%) between square and rectangle patterns. n = number of individual cells measured. * *p*<0.05, *** *p*<0.001, n.s; = not statistically significant.

Most of our analysis focused on the exogenous SMYD3 because of the specificity and efficiency of the anti-Flag antibody, but all experiments with the endogenous protein provided the same conclusions. The total amount of SMYD3 protein was not affected by cell geometry [Figure S2]. We calculate the nuclear:cytoplasmic ratio (expressed as the [Mean intensity per nuclear pixel staining]/ [Mean intensity per cytoplasmic pixel staining]) over a range of patterns areas (500-1800 μm^2^) and aspect ratios: squares (1:1) or rectangles (1:5 or 1:8).

We consistently observed lower nuclear staining for SMYD3 on all rectangles, compared to squares [Figures 1b, S1a and S1d]. Again, the same observations were confirmed whether we measured the exogenous SMYD3-HA-Flag [Figure 1b and Figure S1a] or the endogenous SMYD3 [Figure S1d]. Furthermore, the nuclear:cytoplasmic ratio showed no significant dependence on the spreading areas, for cells plated on square patterns, as well as on rectangular micropatterns. Control experiments with eGFP protein, which freely diffuses through the nuclear pores [^41^], showed that the repartition of a freely diffusing protein is hardly affected by geometry [Figure 1c and Figure S1b]. The difference in the nuclear:cytoplasmic ratio of eGFP between cells plated on square patterns or on rectangle micropatterns (1200 μm^2^ in area) was significant, but very small; less than 3%, as compared to about 30% for SMYD3-HA-Flag and about 65% for endogenous SMYD3. Thus, the nuclear localisation of SMYD3 appears to be influenced by cell geometry, notably mainly by the aspect ratio of the cell.

### Correlating SMYD3 cellular distribution with lysine methylation

The SMYD3 methyltransferase has a number of reported substrates, depending on the cell type and cellular state. These include nuclear histone substrates (e.g. histone H3K4, H4K5, and H2A.Z.1) [^31^, ^32^, ^33^], cytoplasmic proteins (e.g. VEGFR1 receptor and the MAP3K2 signaling kinase) [^34^, ^24^] and interacting proteins (e.g. p53 and HSP90) [^42^,^43^]. Thus, changes in nuclear vs cytoplasmic distribution could likely affect SMYD3 protein interactions and substrate methylation patterns. To investigate whether changes in SMYD3 localisation correlated with lysine methylation, we performed experiments with antibodies recognizing tri-methylated (Kme3) or bi-methylated (Kme2) lysine. We found a strong correlation between the SMYD3-HA-Flag localisation and Kme3 staining, in terms of nuclear:cytoplasmic ratios [Figure 2a]. Furthermore, image analysis suggested a co-localisation of SMYD3 and Kme3 staining [Figure 2b], which was quantitatively confirmed by a high value of the Pearson correlation coefficient, both for cells spread on square micropatterns and on rectangular micropatterns, and both for nuclear and cytoplasmic staining, yet with a better correlation for the cytoplasm [Figure 2c-d]. This correlation was restricted to tri-methylated lysine residues; we failed to observe a correlation when Kme2 antibodies were tested [Figure 2e]; neither in the nucleus (Pearson coefficient~0), nor in the cytoplasm (Pearson coefficient <0.3). Thus, the effect of cell geometry on SMYD3 localisation appeared to directly correlate with lysine tri-methylation in both the nucleus and the cytoplasm.

**Figure 2:**
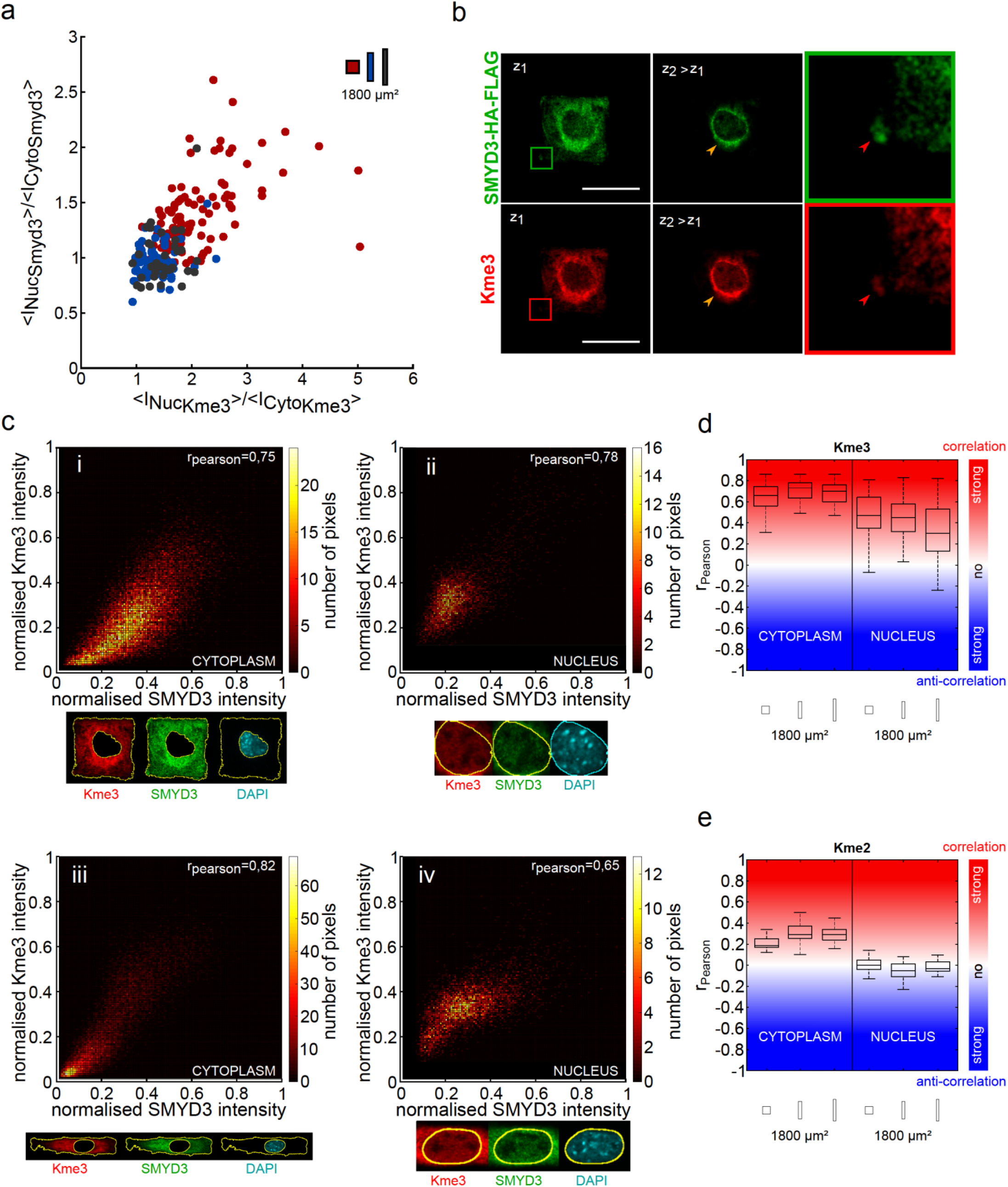
SMYD3 cellular distribution correlates with the lysine trimethylation (Kme3). **(a)** The increased nuclear distribution of SMYD3-HA-FLAG on square patterns (red dots; n=105) correlates with higher nuclear staining for lysine tri-methylation marks (Kme3). Conversely, rectangle patterns (blue and grey spots; n=68 and 37, respectively) have higher SMYD3 and Kme3 in the cytoplasm. **(b)** Confocal microscopy images of a C2C12 cells spread on a square micropatterns, showing the co-localisation of SMYD3-HA-FLAG distribution (green) and lysine tri-methylation Kme3 (red) marks. The magnified square highlights SMYD3 Kme3 colocalisation. Scale bars: 30 μm. **(c)** Upper panels: Detailed representation of the SMYD3 and Kme3 co-localisation within (i) the cytoplasm and (ii) the nucleus for a cell spread on a square micropattern. Lower panels: The same quantification for a cell spread on a rectangle micropattern showing cytoplasmic (iii) and nuclear (iv) quantification. **(d)** Graphical representation of the correlation (Pearson coefficient) between SMYD3 and Kme3 lysine tri-methylation localisation. **(e)** Graphical representation showing a quantified lack of correlation (Pearson coefficient) between SMYD3 and Kme2 lysine di-methylation localization. n = numbers of individual cells measured. Kme3: square n=105, rectangle 1:5 n=68, rectangle 1:8 =37. Kme2: n=24, n=18, n=19.

### The dynamics of SMYD3 nucleo-cytoplasmic shuttling and the role of the cytoskeleton

Nothing is known about the mechanisms underlying SMYD3 localisation and we failed to identify a clear nuclear localization signal (NLS) or nuclear export signal (NES). To investigate nucleo-cytoplasmic shuttling mechanisms, we treated C2C12 cells with Leptomycin B (LMB). Low LMB concentrations bind to CRM1/exportin 1 and block the nuclear export of many proteins [^44^, ^45^, ^46^]. We found that LMB treatment led to nuclear accumulation of SMYD3-HA-Flag in C2C12 cells [Figure 3a]. Although this LMB-induced nuclear accumulation was observed for both geometries, squares and rectangles, it was most marked in cells plated on square than on rectangular micropatterns (~30% vs 20% increase in the nuclear:cytoplasmic ratio) [Figure 3b], which suggests a faster nuclear import for cells plated on squares. To investigate the dynamics of the SMYD3 redistribution, we performed laser bleaching experiments on the cell nucleus or the cytoplasm of cells transfected with SMYD3-eGFP and used fluorescence recovery after photobleaching (FRAP) analysis to follow the nuclear recovery timing [Figure 3c-f]. FRAP analysis experiments showed that the nuclear export of SMYD3-eGFP is faster than its import, which is in agreement with a mainly cytoplasmic localisation. Furthermore, the cell geometry (square vs rectangular micropatterns) impacted the speed of nuclear import, rather than the nuclear export dynamics [Figure 3e-f]. The recovery rate of SMYD3 in the nucleus was significantly higher in square patterns compared to rectangles [Figure 3e], whereas the recovery rates in the cytoplasm were similar, independent of the geometry [Figure 3f].

**Figure 3:**
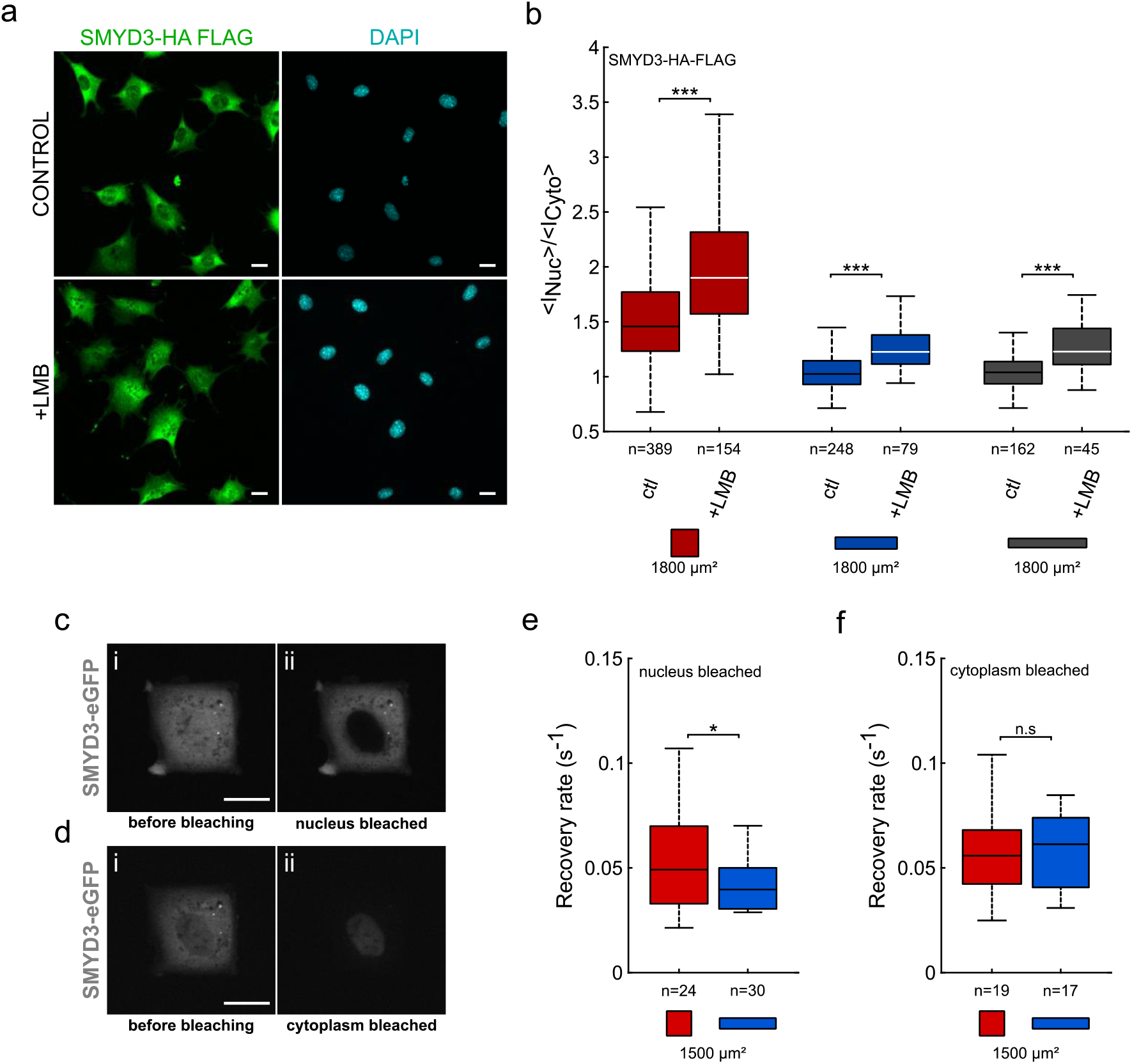
Cell geometry impacts the dynamics of SMYD3 nuclear import. **(a)** Micrographs of C2C12 cells treated with the nuclear export inhibitor leptomycin B (+LMB) causing increased SMYD3 nuclear localisation. **(b)** Quantification of the nuclear:cytoplasmic (<I_Nuc_>/<I_Cyto_>) distribution of SMYD3 in C2C12 cells treated with Leptomycin B (+LMB) compared to untreated controls (ctl) following plating on square (red) or rectangle (1:5, blue) or elongated rectangle (1:8, grey) micropatterns (area=1800 μm^2^). **(c)** FRAP analysis of SMYD3-eGFP localization, before (i) or after (ii) nuclear photo-bleaching. **(d)** FRAP analysis of SMYD3-eGFP localization, before (i) or after (ii) cytoplasmic photo-bleaching. **(e)** Quantification of the recovery rate of fluorescence intensity after nuclear photo-bleaching of C2C12 cells spread on square (red) or rectangle 1:5 (blue) 1500 μm^2^ micropatterns. **(f)** Quantification of the recovery rate of fluorescence intensity after cytoplasmic bleaching of cells spread on square (red) or 1:5 rectangle (blue) 1500 μm^2^ micropatterns. n = number of individual cells measured. * *p*<0.05, *** *p*<0.001, n.s; = not statistically significant.

These drug-treatment and photobleaching experiments clearly showed that nuclear import of SMYD3 is impacted by cell geometry; i.e. relocalisation is faster for cells plated on square patterns than for cells on rectangles. It is well-established that the actin cytoskeleton plays a major role in mechano-sensing phenomena. We therefore treated C2C12 cells with drugs that disrupt the cytoskeleton or impair acto-myosin contractility and measured the effect on SMYD3 distribution. We treated cells with Blebbistatin (that inhibits acto-myosin contractility) or Y-27632 (a pharmacological inhibitor of the Rho-associated protein kinase, ROCK signaling pathway) [Figure 4]. We also treated cells with the Latrunculin A or Latrunculin B inhibitors which disrupt actin filaments. In all cases, perturbing the cellular cytoskeletal integrity increased the nuclear localization of SMYD3-HA-Flag [Figure 4b], on both square and rectangular micropatterns [Figure 4b]. Finally, we tested whether cell geometry impacted the F/G-actin ratio leading to different nuclear localization of SMYD3 Figure S3]. We did not find any correlation between cell shape and the total amount of F-actin, nor between SMYD3 nuclear:cytoplasmic ratios and the amount of F-actin [Figure S3a-d]. Thus, the import of SMYD3 into the nucleus in response to mechano-sensing signals is regulated by the acto-myosin cytoskeleton.

**Figure 4:**
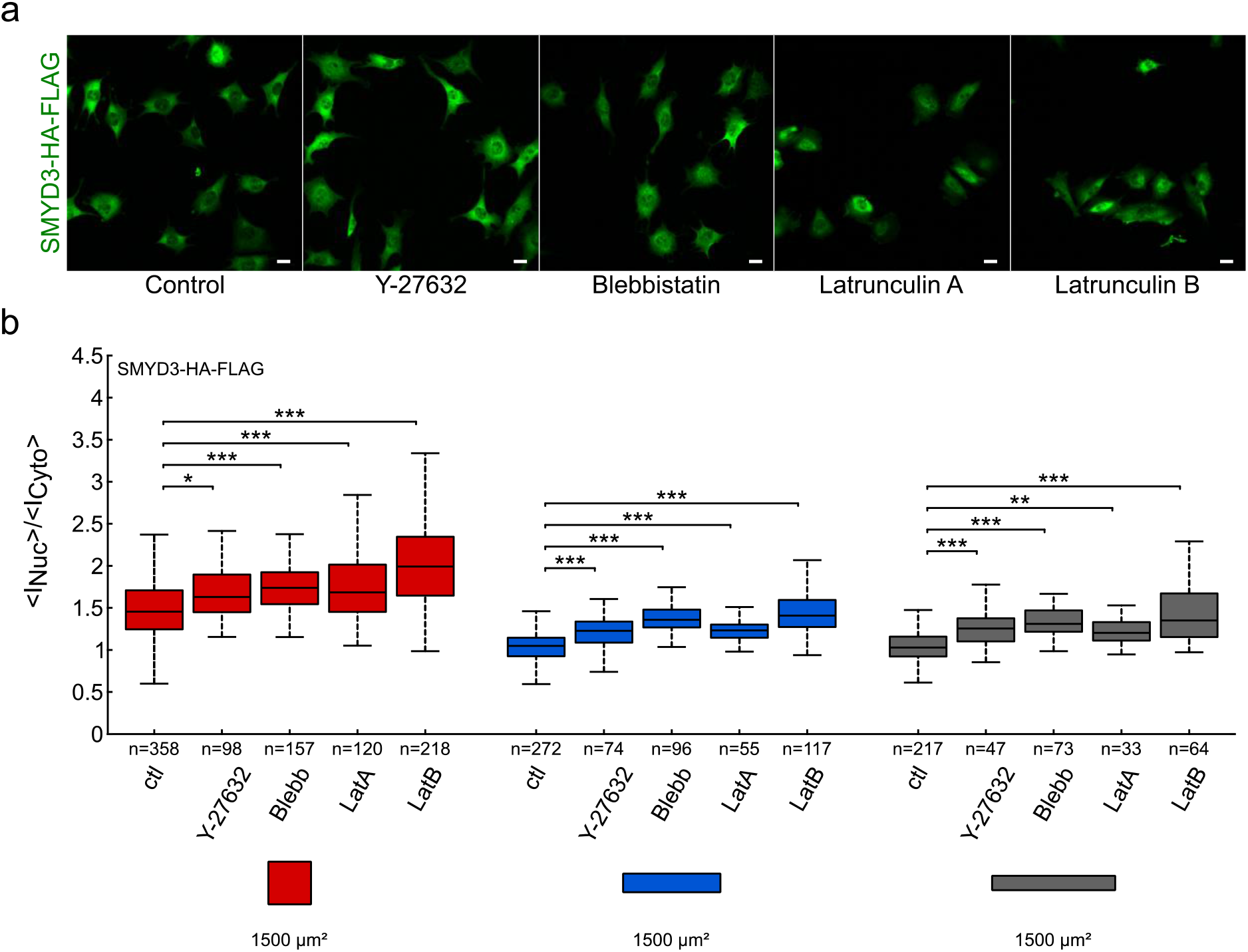
The impact of cytoskeleton disorganisation on SMYD3 cellular distribution. **(a)** Micrographs of SMYD3-HA-FLAG localisation in C2C12 cells treated with drugs targeting cell contractility (Y-27632 or blebbistatin, Blebb) or actin polymerisation (Latrunculin A or Latrunculin B). **(b)** Quantification of increased nuclear localization of SMYD3 upon drug treatment of cells plated on square (red), or rectangle (1:5, blue, 1:8, grey) patterns. Pattern area = 1800 μm^2^. Cells were treated with Y-27632 or blebbistatin (Blebb) or Latrunculin (LatA or LatB). n = number of individual cells measured. * *p*<0.05, ** *p*<0.01, *** *p*<0.001, n.s; = not statistically significant.

### Cell geometry impacts the nucleo:cytosplasmic distribution of a range of regulatory factors

We hypothesized that SMYD3 is not the only transcriptional regulator with a dynamic nuclear-cytoplasmic distribution in response to geometric cues. We sought to test whether other reported mechano-sensitive proteins are also impacted by the square vs rectangle cell geometry. We used our experimental set-up to investigate nuclear:cytoplasmic ratios for C2C12 cells spread on square or rectangular fibronectin over a range of pattern areas (500-1800 μm^2^) [Figure 5 and Figure S4]. We tested localization of another methyltransferase, SETDB1, which was also recently reported to be cytoplasmic [^25^]. We tested G-actin, which is known to play a role in gene transcription [^47^]. We also tested the TAZ protein, a transcriptional regulator that is a paradigm example of mechanical regulation and nucleo-cytoplasmic shuttling [^8^]. We observed in all cases that nuclear:cytoplasmic ratios were higher for cells plated on square micropatterns, compared to rectangular patterns [Figure 5 and Figure S4]. Notably, we found that the TAZ protein was affected by cell size, becoming more nuclear upon spreading on larger and larger areas [Figure 5a-b]. However, we noted that this phenomenon was most remarkable on square patterns,compared to rectangles (aspect ratio 1:5 or 1:8) [Figure 5b and Figure S4b]. We observed the same results when we analysed the nucleo:cytosplasmic distribution of the related YAP protein [Figure S4a]. In contrast, the SETDB1 methyltransferase [Figure S4c] and the G-actin, stained with DNase 1, [Figure 5c-d] distributions behaved similarly to the SMYD3 methyltransferase; the nucleo:cytosplasmic ratios did not significantly depend on spreading area *per se*, but were higher in cells plated on square patterns (in the range 1200-1500μm^2^) [Figure S4c]. Hence, the cell geometry affects the subcellular localization of a range of different proteins, including transcription regulators and epigenetic modulators.

**Figure 5:**
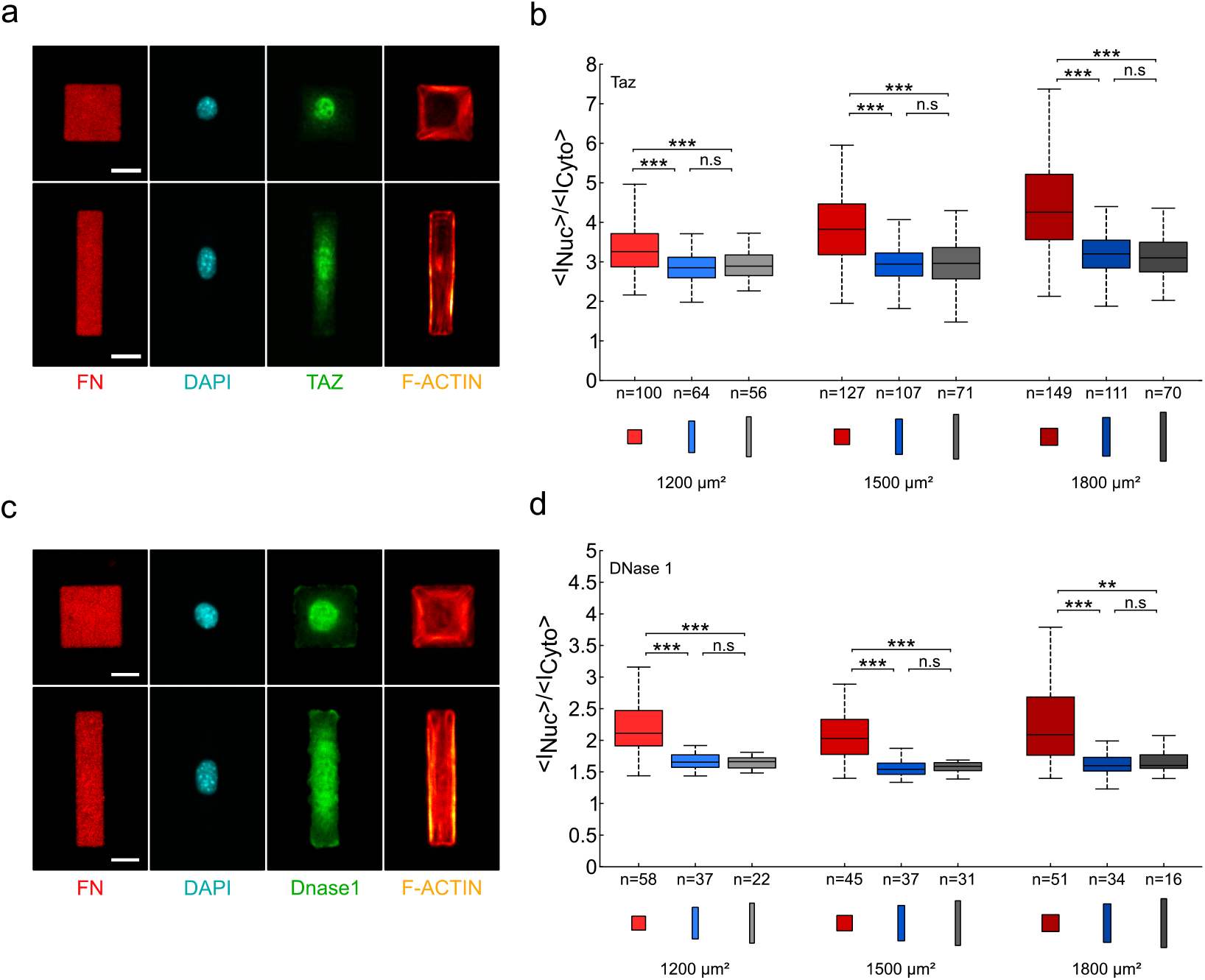
Cell geometry affect the nucleo:cytoplasmic distribution of the TAZ transcription factor and Dnase1. **(a)** Micrographs of TAZ (green) localisation in C2C12 cells spread on square and rectangles (1:5) micropatterns. The micrographs show the fibronectin (FN) patterning, the nuclear DNA staining (DAPI), TAZ (green) location and F-actin (SiR-actin). The TAZ nuclear:cytoplasmic distribution appears different between square and rectangle patterns. Scale bars: 20 μm. **(b)** Quantification of the nuclear:cytoplasmic (<I_Nuc_>/<I_Cyto_>) distribution ratio for Taz over a range of pattern areas (1200-1800 μm^2^) and geometries: squares (1:1 aspect ratio, red), rectangles (1:5 aspect ratio, blue) and elongated rectangles (1:8 aspect ratio, grey). The median value of the ratio is 13 to 37% higher on squares than on rectangles. **(c)** Micrographs of Dnase1 (green) localisation in C2C12 cells spread on square (1:1) or rectangle (1:5) micropatterns. The micrographs show the fibronectin (FN) patterning, the nuclear DNA staining (DAPI), Dnase 1 (green) localisation and F-actin (SiR-actin). The Dnase1 nuclear:cytoplasmic distribution appears different between square and rectangle patterns. Scale bars: 20 μm. **(d)** Quantification of the nuclear:cytoplasmic (<I_Nuc_>/<I_Cyto_>) distribution ratio for Dnase1 over a range of pattern areas (1200-1800 μm^2^) and geometries: squares (1:1 aspect ratio, red), rectangles (1:5 aspect ratio, blue) and elongated rectangles (1:8 aspect ratio, grey). The median value of the ratio is about 30% higher on squares than on rectangle patterns. n = number of individual cells measured. ** *p*<0.01, *** *p*<0.001, n.s; = not statistically significant.

## Discussion

Our study represents a first step towards placing lysine methylation on the pathway from extracellular mechanotransduction to nuclear outcomes. We hypothesized that lysine methylation and chromatin modifiers could offer a chain in the missing conceptual link between transient mechanical signals that are transduced in response to extracellular rigidity and the long-term maintenance of differentiation programs. Lysine methylation events are versatile (lysine can be modified on three different positions me1, me2 or me3, with different functional consequences) and relatively stable. The finding that SMYD proteins have both nuclear and cytoplasmic substrates broadens their potential importance in regulating cellular function in physiological and pathological contexts. We propose, for the first time, that the mechano-sensitivity of SMYD3 distribution could change its cellular substrates and subsequent nuclear vs cytoplasmic functions.

Many previous studies demonstrated that cell spreading impacts nuclear import of transcriptional regulators (e.g. YAP, TAZ, NFκB) [^8^, ^48^], but rarely have these studies carefully examined the relative impact of shape, size or geometry. Our experimental system of fibronectin-coated micropatterns allowed us to control the effect of shape, size and aspect ratio on the localization of different proteins, including transcription regulators such as YAP/TAZ and epigenetic regulators such as SMYD3 or SETDB1. Notably, we observed that increased size led to nuclear accumulation of TAZ and YAP coactivators, as previously reported [Figure 5a-b and Figure S4]. However, we also observed that the nuclear:cytoplasmic ratio of YAP and TAZ depend on geometry, with a preference to nuclear accumulation on squares as compared to rectangles, as recently reported in C2C12 cells [^49^]. Furthermore, the increase in nuclear localization with spreading area is much more marked for squares than for rectangles. Intriguingly, the spatial redistribution of SMYD3 showed a relatively low sensitivity to cell spreading area, but a marked sensitivity to cell shape, with preferred cytoplasmic localization for elongated cells (rectangles with an aspect ratio of 1:5 or 1:8) as compared to isotropically shaped cells (squares). These results highlight a novel distinction between square vs rectangular geometries in regulating a wide range of factors and merits further investigation. The preference for cytoplasmic accumulation on rectangular patterns was clearly evident for other factors tested, e.g. YAP/TAZ and the SETDB1/ESET methyltransferase [Figure 5 and Figure S4] and SMAD proteins (not shown). Notably, cytoplasmic relocalisation of SETDB1 was previously linked to muscle differentiation [^25^]. Furthermore YAP/TAZ has been shown to regulate proliferation [^50^, ^51^, ^52^]. Finally, accumulation of H3K27me3 was observed during mechanically regulated lineage commitment of epidermal progenitor cells [^53^]. It is thus tempting to speculate that transitions of aspect ratios, for example during the myogenic differentiation from isotropically shaped myoblasts to elongated myocytes and fused myotubes, could induce mechanical signals that affect the nuclear:cytoplasmic partitioning of regulatory factors. SMYD3 is linked to muscle differentiation and appears to play a nuclear transcriptional role at earlier time points in myogenesis [^35^]. Furthermore, the deacetylase HDAC3 was reported to be differentially activated by different geometries (aspect ratios) in mouse fibroblasts [^54^]. It would be interesting to look for other differentiation scenarios when methyltransferases switch from nuclear to cytoplasmic functions in response to mechanical signaling.

Our experiments with drug treatments [Figures 3 and 4] suggest that SMYD3 relocalisation is due to mechano-induced import mechanism and that the SMYD3 distribution is regulated by the acto-myosin contractile cytoskeleton. SMYD3 does not have an obvious NLR or NES sequence and further work will be required to define the mechanism of shuttling. An obvious question for future studies is what are the functional consequences of SMYD3 relocalisation on the methylation of cytoplasmic vs nuclear substrates and gene regulation? There is a debate about what are the most biologically relevant SMYD3 histone targets and few gene targets have been identified. It is likely that that substrates and targets are cell-type specific. We previously reported that SMYD3 regulates *Mmp-9* transcription in fibrosarcoma cells [^27^]. This gene seems not be a SMYD3 target in mouse myoblasts (unpublished data), but it is intriguing that the *Mmp-9* gene is regulated by matrix stiffness and mechanotransduction in other systems [^55^]. Also, the activation of cytoplasmic targets is determined by the cellular context, making it difficult to directly predict the mechanistic insights from our experimental system. Our experiments with pan-Lysine-me3 antibodies support a correlation between localization and methyltransferase activity, especially in the cytoplasm, where the correlation is even better than in the nucleus. The relocalisation of SETDB1/ESET has been linked to release from genomic targets and to contribute to muscle differentiation [^25^]. We favour a model in which specific cell geometries affect nuclear vs cytoplasmic repartitioning of SMYD3 (and other methyltransferases) leading to shifts in substrate encounters and regulation by methylation.

## Methods

### Stamp fabrication

The desired shapes (rectangles and squares) were designed using L-Edit software. A chromium mask was used to create a silicon/SU8 mould with rectangular and square shapes. It was generated by a photolithographic process. The mould was used to carry polydimethylsiloxane (PDMS) overnight at 60°C, the PDMS layer was peeled off and washed on an ultrasonic bath containing a solution of 70% of ethanol for 15 minutes.

### Microcontact printing

The structured PDMS stamp was dried with clean dry air and inked for 45 min at room temperature (RT) with a mixture of fibronectin solution from bovine plasma (FN) (Sigma Aldrich) and labeled (cy3.5 Invitrogen labeling kit) fibronectin from human plasma (Roche), the final concentration used for the experiments was 50 μg/ml (containing 2/3 of bovine and 1/3 of human FN). A thin layer of PDMS was exposed to UV/O3 irradiation for 7 min. The FN-coated stamp was washed with phosphate buffer saline (PBS), dried with clean dry air and then put on contact onto the PDMS layer surface for few seconds. The surface was passivated for 45 min with a 0.2% solution of Pluronic F-127.

### Cells culture and drug treatment

C2C12 myoblasts cells were cultured in DMEM (high glucose, pyruvate, glutamax) medium (FBS 15%) and 1% pen-strep. 5000 cells /cm^2^ were plated on the microprinted PDMS layers for 3h at 37°C in 5% CO2. C2C12 stably expressing HA-Flag-SMYD3 and C2C12 transfected (Lipofectamine) with eGFP were used. The chosen drugs were added in the media at the indicated low concentrations to allow spreading and to destabilize the cytoskeleton. The indicated times are measured from the end of the experiment (total time of experiment 3h): Rock inhibitor 10 μM 1h (Cell Guidance Systems), Blebbistatin 5 μM 1h (Sigma Aldrich), Latrunculin A 0.12 μM 2h45 (Sigma Aldrich), Latrunculin B 80 nM 30min (Sigma Aldrich), and LMB 20 nM 2h (Sigma Aldrich).

### Construction of C2C12 cell line stably expressing HA-Flag-SMYD3

SMYD3 cDNA was amplified by PCR using the following primer pair: (for) cgatgctctcagtgccgcgt and (rev) ggatgctctgatgttggcgtc. cDNA was cloned into the pREV retroviral vector, kindly obtained from the laboratory of Dr S. Ait-Si-Ali (CNRS, UMR7216, Paris). The plasmid contains an epitope tag (3 HA- and 3 Flag-tag) in 5’ of the cloning site and a selection marker. C2C12 myoblasts stably expressing double-tagged Flag-HA-SMYD3 protein were established using retroviral transduction strategy as previously described [^56^]. C2C12 cells stably integrating pREV-SMYD3 or pREV control vector were sub-cloned to obtain 100% positive clonal populations. Stable ectopic SMYD3 expression in these clones was validated by Western blot and immunofluorescence analysis, using anti-HA and anti-FLAG antibodies. We selected two C2C12 positive clonal populations for our further experiments.

### Immunochemistry and labeling

After 3h in culture on the microprinted PDMS layers, cells were fixed with 4% PFA at room temperature (RT) for 15 min and then permeabilized with 0.2% Triton X-100 in PBS for 5 min at RT and washed 3 times in PBS. The immunochemistry process was performed following these steps: the fixed cells were incubated for 30 min in a blocking solution (0.2% tween, 1% Bovine Serum Albumine BSA, 1% Foetal Bovine Serum FBS) at RT, then the primary antibody was incubated for 40 min, washed three times in PBS-tween 0.2%, the secondary was incubated for 30 min, washed three times with PBS. The nucleus was then labeled with 4’,6-diamidino-2-phenylindole (DAPI) for 30 min at RT and the actin filaments (F-actin) with SiR-actin (Spirochrome) at 100 nM in PBS overnight at 4°C. The antibodies used for the immunochemistry were: anti-Flag (Sigma Aldrich) 1/1000, anti-SMYD3 (Abcam) 1/1000, anti-TAZ (Cell Signaling Technology) 1/500, anti-YAP (Santa Cruz Biotechnology) 1/250, anti-Pan lysine trimethylated me3 (Cell Signaling) 1/1000, anti-Pan lysine dimethylated me2 (Cell Signaling) 1/1000, anti-SMYD3 (Abcam) 1/300, DNase 1 Alexa Fluor 594 (Life Technologies) 1/1000, anti-SETDB1 (Santa Cruz Biotechnology) 1/200.

### Image analysis

Homemade codes were written to analyze experimental data, using Matlab and ImageJ software and Miji plugin. Using this software, we developed an automatized pipeline to segment and measure many parameters such as intensity, shape, aspect ratios, nuclear to cytoplasmic ratios of cells on micropatterns.

### Colocalisation experiments

Images have been acquired using a spinning disk confocal microscope with a 60X objective (Olympus/Yokogawa/Andor). For each cell, a characteristic plane (XY) of each channel has been used to plot the intensity of Kme3 (or Kme2) vs SMYD3 intensity. This has been done for the nucleus and the cytoplasm separately. The estimation of the correlation of the intensities has been done using the Pearson’s correlation coefficient 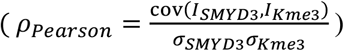.

### FRAP experiments

Cells transfected with SMYD3-eGFP were plated on squares and rectangles micropatterns (area of 1500 μm^2^). To measure the nuclear import (export), we bleached the whole nucleus (cytoplasm) and we measured the fluorescence recovery after the photobleaching in the nucleus. Then, the experimental curves have been fitted using exponential functions.

### Statistical analysis

Different statistical analyses were performed depending on the number N of distributions and on the number n of measurements in each distribution. When n≥30, the significance was tested using a Student test for the comparison between N=2 distributions and an Anova one way test (Tukey-Kramer posthoc test) when N>2. When n<30, the normality of the distributions was first tested using an Anderson-Darling test. For normal distributions, the tests were the same as when n≥30. If the distributions were not normal, the significance was tested using a Wilcoxon rank sum test for the comparison between N=2 distributions and a Kruskal-Wallis test (Dunn and Sidak posthoc test) when N>2.

## Author Contribution Statement

SH and JBW developed the conceptual framework, sought funding and provided overall supervision.

DP and AR performed the experiments. DP performed all the quantitative analysis of the results.

DP, SM, SH and JBW designed the study, analysed the findings and wrote the manuscript.

## Acknowledgements

We thank members of the Hénon and Weitzman laboratories and members of the UMR7216 Epigenetics and Cell Fate for discussions and advice on this study. We acknowledge the ImagoSeine core facility of the Institut Jacques Monod (member of the France BioImaging, ANR-10-INBS-04). This work was supported by the LabEx “Who Am I?” #ANR-11-LABX-0071 and the Université de Paris IdEx #ANR-18-IDEX-0001 funded by the French Government through its “Investments for the Future” program. Additional funding from AFMTELETHON (#16146), and the Fondation ARC pour la Recherche sur le Cancer (ARC n°155029). JBW is a Senior member of the Institut Universitaire de France (IUF) and SM is a Junior member (2012ND 3369).

