## Supplementary material for "The impact of cell geometry and the cytoskeleton on the nucleo-cytoplasmic localisation of the SMYD3 methyltransferase suggests that the epigenetic machinery is mechanosensitive": hh

### Supplementary Information

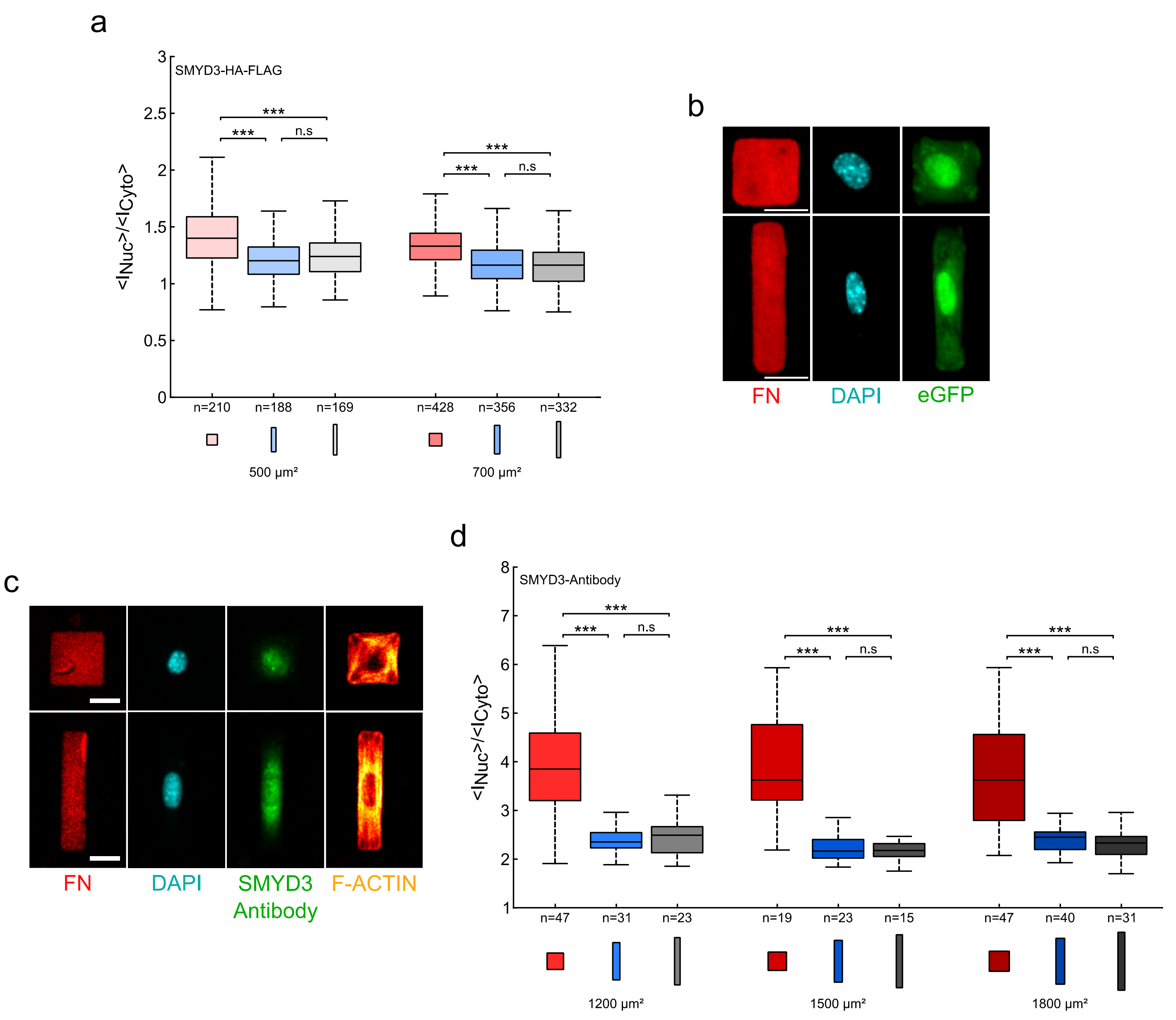


**Figure S1**: **Cell geometry on fibronectin micropatterns regulates exogenous and endogenous SMYD3 distribution.**

**a)** Quantification of the nuclear:cytoplasmic (<$I_{Nuc}$>/<$I_{Cyto}$>) distribution ratio for exogenous SMYD3-HA-Flag on small pattern areas (500 and 700 µm²) and geometries: squares (1:1 aspect ratio, red), rectangles (1:5 aspect ratio, blue) and elongated rectangles (1:8 aspect ratio, grey). The median value of the ratio is about 15% higher on squares than on rectangle patterns.

**b)** Micrographs show the fibronectin (FN) patterning, the nuclear DNA staining (DAPI), and localization of control eGFP in C2C12 cells plated on square or rectangle patterns (1200 µm²). See Figure 1c for quantification data.

**c)** Non-transfected C2C12 cells were plated on different fibronectin micropatterns with the same area (1200 µm²), but different aspect ratios; square (1:1, upper panels) or rectangle (1:5, lower panels). The micrographs show the fibronectin (FN) patterning, the nuclear DNA staining (DAPI), endogenous SMYD3 protein detected with a specific antibody (SMYD3) and F-actin.

**d)** Quantification of the nuclear:cytoplasmic (<$I_{Nuc}$>/<$I_{Cyto}$>) distribution ratio for endogenous SMYD3 protein over a range of pattern areas (1200-1800 µm²) and geometries: squares (1:1 aspect ratio, red), rectangles (1:5 aspect ratio, blue) and elongated rectangles (1:8 aspect ratio, grey). median value of the ratio is 48 to 67% higher on squares than on rectangle patterns.

n = number of individual cells measured. *** $p$<0.001, n.s; = not statistically significant.

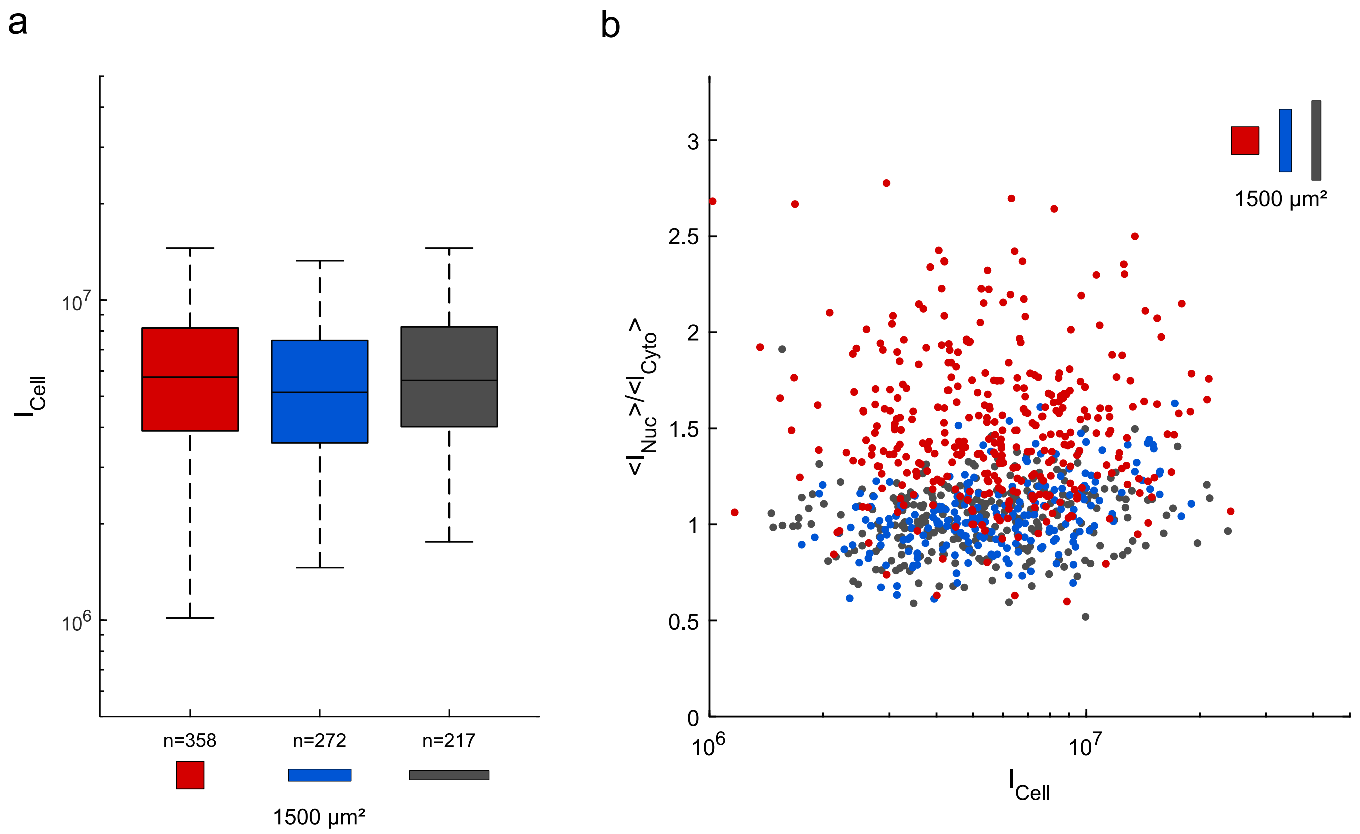


**Figure S2**: **Cell geometry does not affect the total intensity of SMYD3**

**a)** Quantification of the total cell intensity of exogenous SMYD3-HA-Flag on pattern area (1500 $\mu m^2$) of different geometries: squares (1:1 aspect ratio, red), rectangles (1:5 aspect ratio, blue) and elongated rectangles (1:8 aspect ratio, grey). n = number of individual cells measured.

**b)** Quantification of the nuclear:cytoplasmic ($<I_{Nuc}>/<I_{Cyto}>$) distribution ratio for exogenous SMYD3-HA-Flag on pattern area (1500 $\mu m^2$) compare to the total intensity for each cell.

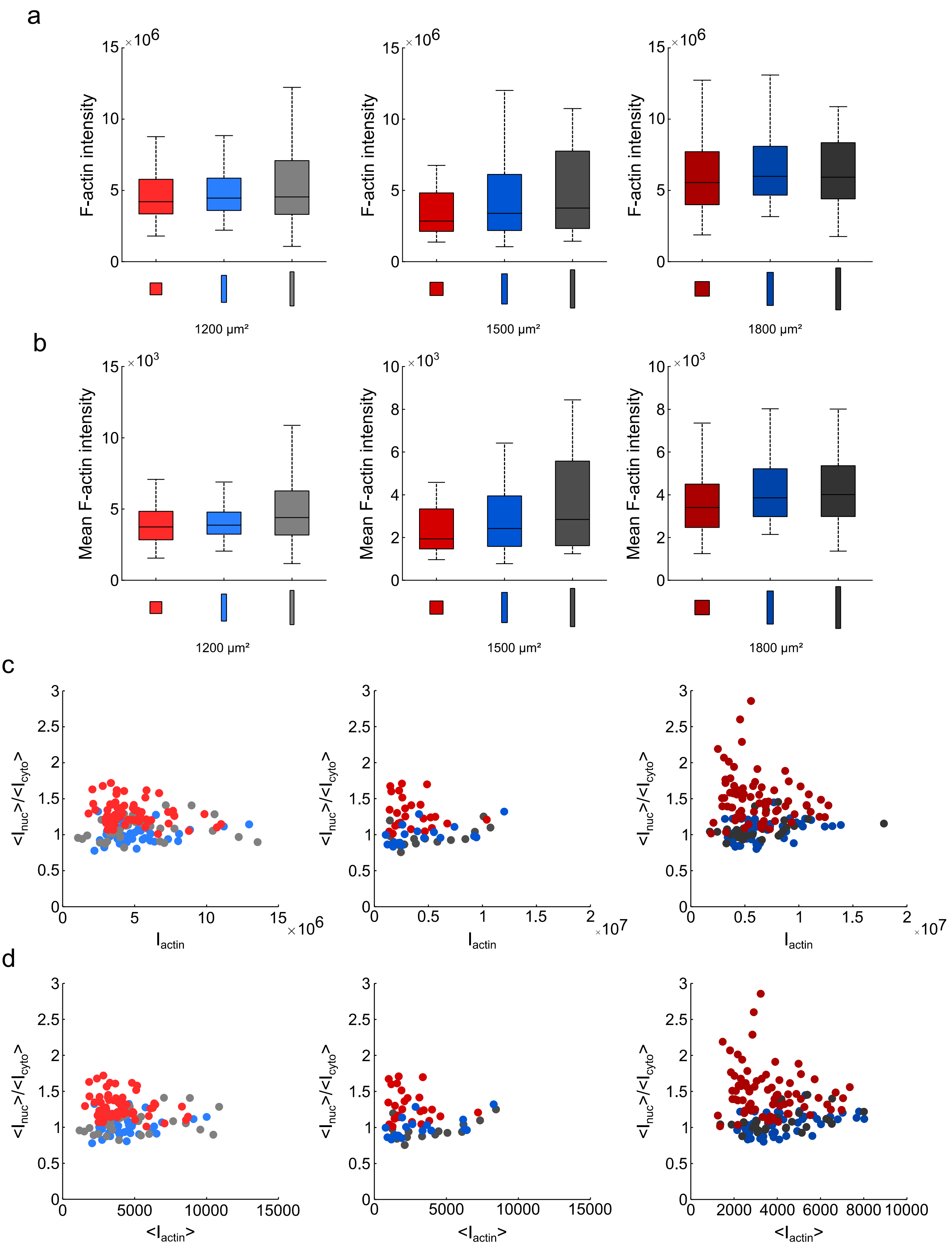


**Figure S3**: **The amount of F-actin in cells spread on micropatterns is independent of cell shape or the SMYD3 distribution.**

- **a)** The total amount of F-actin, quantified by the intensity of SiRactin staining for cells plated on micropatterns with different areas (1200 to 1800 µm²) and shapes: squares (1:1 aspect ratio, red), rectangles (1:5 aspect ratio, blue) and elongated rectangles (1:8 aspect ratio, grey).
- **b)** The mean amount of F-actin per pixel, under the same conditions.
- **c)** Quantification of the nuclear:cytoplasmic ($<I_{Nuc}>/<I_{Cyto}>$) distribution ratio for exogenous SMYD3-HA-Flag on pattern area (1500 µm²) compare to the total intensity for each cell.
- **d)** Quantification of the nuclear:cytoplasmic ($<I_{Nuc}>/<I_{Cyto}>$) distribution ratio for exogenous SMYD3-HA-Flag on pattern area (1500 µm²) compare to the total mean intensity of the SiRactin channel for each cell.

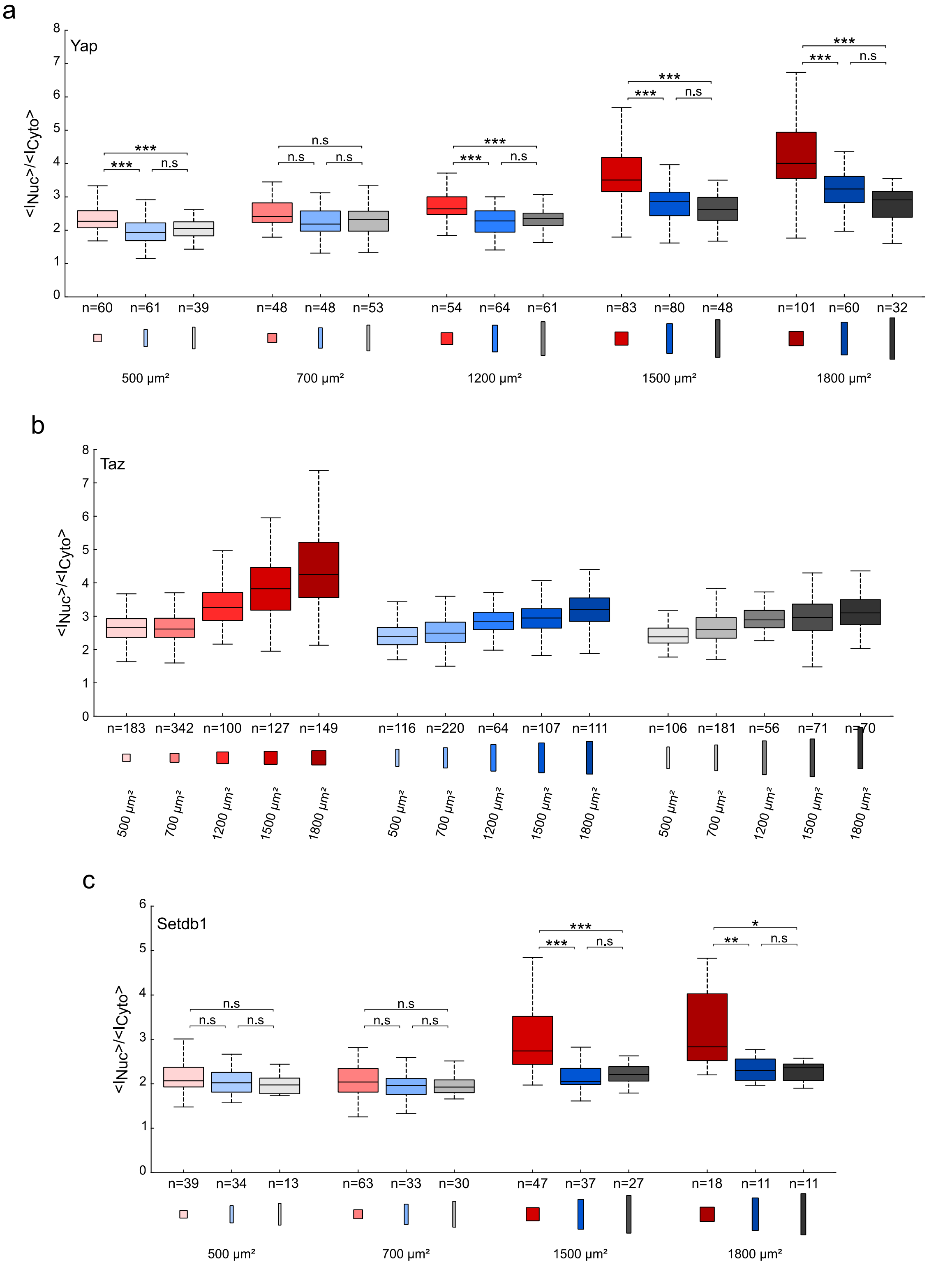


**Figure S4**: **Cell geometry on fibronectin micropatterns regulates the localization of a range of regulatory factors.**

**a)** Quantification of the nuclear:cytoplasmic ($<I_{Nuc}>/<I_{Cyto}>$) distribution ratio for endogenous Yap protein in cells plated on different pattern areas (500-1800 µm²) or shapes: squares (1:1 aspect ratio, red), rectangles (1:5 aspect ratio, blue) and elongated rectangles (1:8 aspect ratio, grey). The median value of the ratio is 10 to 24% higher on squares than on rectangles.

**b)** Quantification of the nuclear:cytoplasmic ($<I_{Nuc}>/<I_{Cyto}>$) distribution ratio for endogenous Taz protein in cells plated on different pattern areas (500-1800 µm²) and shapes: squares (1:1 aspect ratio, red), rectangles (1:5 aspect ratio, blue) and elongated rectangles (1:8 aspect ratio, grey).

**c)** Quantification of the nuclear:cytoplasmic ($<I_{Nuc}>/<I_{Cyto}>$) distribution ratio for endogenous Setdb1 protein in cells plated on different pattern areas (500-1800 µm²) and geometries: squares (1:1 aspect ratio, red), rectangles (1:5 aspect ratio, blue) and elongated rectangles (1:8 aspect ratio, grey). For areas 1500 and 1800 µm², the median value of the ratio is 20-33% higher on squares than on rectangles.

n = number of individual cells measured. * p<0.05, ** *p*<0.01, *** *p*<0.001, n.s; = not statistically significant.
